# Jasmonic acid signalling is targeted by a smut fungal Tin2-fold effector

**DOI:** 10.1101/2024.07.08.602457

**Authors:** Summia Gul, Gabriel Mendoza-Rojas, Natascha Heßler, Stefanie Galle, Sander H.J. Smits, Florian Altegoer, Vera Göhre

## Abstract

In plants, jasmonate signaling is a hub integrating environmental cues with growth and development. Due to its role in balancing defense responses against pathogens, it is a target of effector proteins from various pathogens. Here, we characterized the fungal effector protein Tue1 from the Brassicaceae smut fungus *Thecaphora thlaspeos*. *T. thlaspeos* naturally infects *Arabis hirsuta* but can also colonize the non-host *Arabidopsis thaliana*. In planta, the fungal protein Tue1 hijacks the plant importin-α dependent nuclear transporter to reach the plant nucleus. It interacts with jasmonate ZIM domain 10 (JAZ10) proteins of both *A. thaliana* and *Ar. hirsuta*. Structure-guided analysis of Tue1 suggests that it binds the Jas motif of JAZ10 indicating a role in stabilization or binding competition with proteins like MYC3 and COI1. A subset of jasmonate-responsive genes is differentially regulated during *T. thlaspeos* infection, proposing a link of the Tue1 function to infection. Tue1 share structural similarity to the Tin2-fold family recently described in the corn smut *Ustilago maydis*. Our study therefore suggests that this structural effector family is expanded across fungal pathogens, although future studies have to reveal whether targeting JAZ-repressors is a conserved mechanism or specifically acquired as an adaptation to its perennial host.

## Introduction

During the evolution of plants, fungi played a critical role in shaping the development of land plants (Hoeksema *et al*., 2018). Most vascular plants are engaged in tight associations with mycorrhizal fungi representing an important aspect of plant terrestrialization (Puginier *et al*., 2022). Not only beneficial but also parasitic interactions of fungi with plants have established, resulting in numerous fungal diseases that strongly influenced ecosystems and even lead to extinctions of plant and animal species (Fisher *et al*., 2012). To shape these interactions, fungi use specialized secreted proteins termed effectors that facilitate the colonization process and e.g. suppress plant immune responses or manipulate central cellular processes of the host (Lanver *et al*., 2017; Lo Presti *et al*., 2015). As many of these proteins evolved in the course of a specific fungal-plant interaction, they often share little sequence homology and lack domains of known function. Although effector proteins are the primary focus of research in the field of plant pathology, biochemical- and studies focusing on structure-function relationships are still critically lacking and the functions of most virulence factors are still poorly understood.

A large group of fungal pathogens are smut fungi with more than 550 Basidiomycete species. These fungi have a narrow host range and infect important crops such as maize, wheat, rice or barley (Zuo *et al*., 2019) peanut (Paredes *et al*., 2024) and potato (Health *et al*., 2018). Smut fungi are biotrophic pathogens that rely on viability of their infected hosts and establish an intimate interaction. While smut species colonizing monocot hosts are best characterized due to their agronomic impact, the dicot smut fungi in particular from the *Thecaphora* clade (Vanky, 2008) are understudied. For example, *Thecaphora thlaspeos* naturally occurs in >15 Brassicaceae host plants including *Arabis* species (Vanky, 2008) and it can colonize the model plant *Arabidopsis thaliana* (Frantzeskakis *et al*., 2017). In recent years, *T. thlaspeos* has been established as a dicot smut model system that can also be genetically modified (Plücker *et al*., 2021). Its genome is a typical smut genome of small size (Courville *et al*., 2019), but the effector repertoire shows some remarkable differences concerning the core effectors of grass smut fungi (Schuster *et al*., 2018).

Effector genes of smut fungi are often but not exclusively organized in clusters that encode small protein families (Kämper *et al*., 2006). Some of these families have been demonstrated to contribute to virulence, while for others no influence on the plant colonization process was observed in lab infections (Kämper *et al*., 2006; Schirawski *et al*., 2010). The sequence variability between these effector genes is high and only a small percentage is conserved among related smut species (Laurie *et al*., 2012). The dicot-infecting smut fungi such as *Melanopsichium pennsylvanicum* and *Thecaphora thlaspeos* even share a smaller proportion of conserved effectors with their grass-infecting relatives (Courville *et al*., 2019; Sharma *et al*., 2014). Despite a close relation to *U. maydis* and *T. thlaspeos* on the genomic level,(Courville *et al*., 2019) the smut fungus *Pseudozyma flocculosa* (*Anthracocystis flocculosa*) that has so far been described as endophyte and biocontrol agent rather than a parasite, even lost specific subsets of effector proteins (Lefebvre *et al*., 2013). By contrast, core effectors have been identified that are functionally conserved among dicot and monocot smuts, such as Pep1 (Courville *et al*., 2019; Hemetsberger *et al*., 2015). Hence, effectors can be grouped into core effectors with shared functions between smut fungi and specific effectors used by individual or small subgroups to support infection.

Previously, we have identified a subset of effector candidates unique to *T. thlaspeos*. Of those, the ***T****hecaphora*-**u**nique **e**ffector 1 (Tue1) caused a strong growth defect and promoted bacterial virulence upon heterologous expression *in planta* (Courville *et al*., 2019) and has therefore turned out as a promising candidate for further investigation. Here, we now provide a detailed characterization of Tue1 and give insights into its molecular function in interfering with plant hormone signalling. Our structure-guided approach reveals that Tue1 belongs to an expanded family of smut fungal effectors that do not share sequence similarities, but have a common structural core. While the characterization of Tue1 in the infection biology of Brassicaceae gives insight into how a perennial pathogen modifies the plant during the extended biotrophic phase, in the future this core structure might be used as a tool to modify plant hormone signalling.

## Materials and Methods

### Accession numbers and sequence analysis

*A. thaliana* sequences were taken from TAIR: *At*JAZ10= At5g13220.1, AtMYC3= At5g46760. *T. thlaspeos* sequences for Tue1 (THTG_04687) and Tue16 (THTG_04669) and *Ar. hirsuta* sequences for JAZ10 (ArH_00064583-RA) were taken from our genome and transcriptome sequencing (Courville *et al*., 2019). Nuclear localization sequences were predicted using NLStradamus (Nguyen Ba *et al*., 2009), Localizer (Sperschneider *et al*., 2017) and cNLSmapper (Kosugi *et al*., 2009).

### DNA amplification and molecular cloning

*tue1* and *tue16* were amplified from cDNA of *T. thlaspeos* strain LF1 excluding the signal peptide (Courville *et al*., 2019). *AtMYC3* (5-242) and *AtJAZ10* were amplified from cDNA of *A. thaliana* Col-0 and *AhJAZ10* was amplified from cDNA of *Ar. hirsuta* RH2015. Expression constructs for the effectors under the control of the 35S promoter in *Nicotiana benthamiana* and *A. thaliana* were generated by Standard GreenGate cloning (Lampropoulos *et al*., 2013). For bimolecular fluorescence complementation assay (BiFC), *AtMYC3*(5-242)*, tue16*, *tue1* and the truncated version were tagged with N-terminal split mVenus and *AtJAZ10* and *AhJAZ10* were tagged with the C-terminal split mVenus. For yeast-two-hybrid, *tue1* lacking the signal peptide was inserted into pGILDA (Clontech) to generate a fusion with the *lexA* gene. Plant targets identified in the yeast-two hybrid screen were cloned as translational fusions with the activation domain in pB42AD (Clontech) for verification. A standard GoldenGate cloning procedure (Zweng *et al*., 2023) was used to generate expression constructs in pEMGB1 backbone*. tue1*, *tue16* and *AtJAZ10* were N-terminally tagged with hexa-histidine. The solubility tag GB1 was fused to the N-terminus of JAZ10.

### Strains and growth conditions

The *E. coli* strain Top10 (Thermo Fisher Scientific) was used for cloning purposes. The *E. coli* strain BL21 (DE3) (Novagen) was used to express all produced proteins in this study. *E. coli* strains were grown at 37 °C in dYT media (tryptone 1.6% (w/v), yeast extract 1% (w/v) and NaCl 0.5% (w/v). The *Agrobacterium tumefaciens* strain C58pMP90 (Koncz & Schell, 1986) was employed for expression of effectors in *A. thaliana* and *N. benthamiana*. *Ag. tumefaciens* strains were grown at 28 °C in dYT media containing the appropriate antibiotics (100 µg/ml spectinomycin, 10 µg/ml rifampicin, 50 µg/ml gentamycin and 10 µg/ml tetracycline)*. Saccharomyces cerevisiae* strain EGY48 (Matchmaker 3 system, Clontech) was used to perform yeast two-hybrid library screen. *S. cerevisiae* strains were grown at 30 °C in YPDA (yeast extract 1.25 % (w/v), peptone 2.5 % (w/v), glucose 2 % (w/v), 0.008 % adenine) and SD-dropout medium (amino acid mix 0.5 % (w/v), yeast nitrogen base 1.7 % (w/v), glucose 2 % (w/v)) at 200 rpm.

### Transgenic *A. thaliana* lines and transient expression in *N. benthamiana*

Transgenic *A. thaliana* Col-0 lines carrying Tue1-Gfp, free GFP and NLS-mCherry constructs were generated via floral dip method (Deak *et al*., 1986). Transgenic seeds were selected on kanamycin, and verified for expression of the respective fluorescent proteins. *N. benthamiana* was infiltrated by *Ag. tumefaciens* for transient expression of Tue1-Gfp, Tue1ΔNLS-Gfp, free Gfp and NLS-mCherry. Protein localization was assessed in the stable lines or in *N. benthamiana* at 3 days post infiltration with Zeiss LSM 880 confocal microscope.

### Protein production and purification

Protein purification was performed as previously described (Zweng *et al*., 2023). Briefly, *E. coli* BL21 (DE3) strains (Novagen) containing pET22b-Tue1-His were grown to an OD600 of 0.6 at 30 °C and protein expression was induces with 0,5 mM IPTG at 20 °C for 20 h. The cells were disrupted through a microfluidizer (M110-L, Microfluidics). Debris-free supernatant was loaded onto Ni-NTA FF-HisTrap columns (GE Healthcare) for affinity purification via the hexahistidine tag. The eluted protein was concentrated using Amicon Ultra-10K centrifugal filters and subjected to size-exclusion chromatography (SEC) using a Superdex S200 Increase 16/600 column. The peak fractions were analyzed using a standard SDS-PAGE protocol, pooled, and concentrated with Amicon Ultra-10K centrifugal filters.

### Selenomethionine incorporation for anomalous diffraction

*E. coli* BL21 (DE3) strains (Novagen) containing pET22b-Tue1-His cultures were inoculated in M9 medium (37.25 g/l Na2HPO4, 16.5 g/l KH2PO4, 2.75 g/l NaCl, 5.5 g/l NH4Cl, pH 7.5) to an OD600 of 0.1 infused with sterile and freshly made SolX solution (1 g/l L-lysine, 1 g/l L-threonine, 1 g/l L-phenylalanine, 0.5 g/l L-leucine, 0.5 g/l L-isoleucine, 0.5 g/l valine, 0.25 g/l selenomethionine, 80 g/l glucose, 100 mM MgCl2, 10 mM CaCl2) and grown to OD600 of 0.6. Protein production was induced with 1 mM IPTG for 20-22 h at 37°C. The cells were harvested and stored at -80 °C or immediately used for protein purification.

### Protein crystallization

Crystallization was performed using MRC-3, 96-well sitting drop plates, and commercial crystallization screening kits at 12 °C. 0.1 µL homogeneous protein solution was mixed with 0.1 µL reservoir solution and equilibrated against 40 µL of the reservoir. After one-week, initial rod-shaped crystals were found which were further optimized by slightly varying the precipitant concentrations. Optimization was also conducted in sitting drop plates (24-well) at 12°C but by mixing 1 µL protein solution with 1 µL of the reservoir solution, equilibrated against 300 µL reservoir solution. Native Tue1-6H crystallized at 12 mg/ml concentration within 2 weeks in 0.05 M NaCl, 1.2M K/Na tartrate, 0.1M imidazole pH 8. Se-Met Tue1-6H crystallized at 12 mg/ml concentration within 1 month in the same condition. Before harvesting the crystals, crystal-containing drops were overlaid with 2 µL mineral oil and immediately flash-frozen in liquid nitrogen.

### Data collection, processing and structure refinement

The data were collected under cryogenic conditions at the EMBL beamline P13 (Deutsches Elektronen Synchrotron; DESY). The data were integrated and scaled using XDS and merged with XSCALE (Kabsch, 2010). The structure of Tue1 was phased by isomorphous replacement using data obtained from single-wavelength anomalous dispersion gathered by incorporating selenomethionine. The structure was manually built in COOT (Emsley *et al*., 2004), and refined with PHENIX (Adams *et al*., 2010). The figures were prepared with ChimeraX (Pettersen *et al*., 2021). All residues were found within the preferred and additionally allowed regions of the Ramachandran plot. Detailed data collection and refinement statistics are listed in the supplementary table 1.

### Yeast two-hybrid assays

*S. cerevisiae* strain EGY48 was transformed with Tue1-LexA-pGILDA (Clontech, MATCHMAKER LexA Two-Hybrid System) and tested for auto-activity. A cDNA library from stress-induced *A. thaliana* in pB42AD (Matiolli and Melotto, 2018) was co-transformed with Tue1-LexA-pGILDA for screening according to the Clontech manual. For the library transformation, the protocol was scaled up to 2400 μl competent cells and 10 μg plasmid DNA of library was used (Matiolli and Melotto, 2018).

Plasmids from interaction candidates were extracted according the manual (Clontech). The selected prey candidates were sequenced and identified by comparison to the *A. thaliana* genome. Full-length gene models for the candidates were obtained from TAIR, and *Ar. hirsuta* homologs were identified from our preliminary *Ar. hirsuta* sequences (Courville *et al*., 2019).

Protein extraction from yeast cells expressing Tue1, *At*JAZ10 and *Ah*JAZ10 was done according to the manual (Clonetech) for Western Blot verification of protein enrichment.

### Bimolecular fluorescence complementation (BiFC)

Translation fusions of the effector candidates and plant targets (*AtMYC3*, *tue16, tue1,* and its truncated versions tagged with with N-terminal split mVenus; *AtJAZ10* and *AhJAZ10* with C-terminal split mVenus) as well as an *NLS-mCherry* construct in *Ag. tumefaciens* strain C58 pMP90 pSOUP were infiltrated into *N. benthamiana* leaves. Signal reconstitution of the split fluorophore was observed by confocal microscopy at 2-3 days post infiltration on a Zeiss LSM 880 microscope. Fluorescence signals were quantified using ImageJ (http://rsb.info.nih.gov/ij) and analysed by mean fluorescence intensity (MFI) and JACoP BIOP plug-in (https://github.com/BIOP/ijp-jacop-b) for the total fluorescent intensity and co-localized signal intensity with NLS marker, respectively.

### Microscale thermophoresis (MST)

Recombinant 6HN-GST-MYC3 (5-242), 6HN-GST-Tue1 and 6HN-GST-Tue16 protein were labeled with Alexa Fluor® 488 (NHS Ester, Lumiprobe GmbH, Germany). Recombinant 6HN-JAZ10 (3,1nM-100 μM, 16 dilution series) was incubated with each labeled protein at 200 nM in 0.01% Tween-20-SEC buffer. 10 μL sample was transferred into a glass capillary (NanoTemper, Munich, Germany) and thermophoresis was detected with NanoTemper Monolith Instrument (NT.115, Munich, Germany), performed with an excitation power of 20% for 30 s and MST power of 40% at an ambient temperature of ∼24 °C. Triplicates of the same dilution were measured. The results were further analyzed by MO Affinity Analysis Software (NanoTemper, Munich, Germany).

## Results

### Tue1 localizes to the plant nucleus

In our previous study, we could show that Tue1 strongly influences the growth of *A. thaliana* upon heterologous expression resulting in significantly smaller rosettes (Courville *et al*., 2019). We were therefore interested to reveal the mechanism underlying this strong phenotype. While Tue1 does not contain any predicted functional domains (Courville *et al*., 2019), Localizer (Sperschneider *et al*., 2017) and cNLSmapper (Kosugi *et al*., 2009) suggested the presence of a nuclear localization signal (NLS) between amino acid 80 and 90 in addition to the predicted signal peptide at the N-terminus. (**Fig. 1A**). To confirm the predicted nuclear localization, we performed heterologous expression *in planta* using GFP-fusions lacking the N-terminal signal peptide in comparison to NLS-mCherry (NLS of At4g19150/N7, (Cutler *et al*., 2000)). Transient expression in *N. benthamiana* showed a clear nuclear localization of Tue1 (**Fig. 1B**). To further validate that the predicted NLS confers the nuclear localization, we generated a construct of Tue1 lacking the NLS (GGVA**KR**P**R**IS). This deletion resulted in a cytoplasmic localization (**Fig. 1C**) suggesting that Tue1 is indeed targeted to the nucleus via this identified sequence stretch. Tue1-Gfp and its truncated variant were expressed *in planta* as full-length protein with no visible degradation (**Fig. 1D**). To investigate whether this fungal NLS can be recognized by the plant nuclear import machinery, we predicted the structure of *A. thaliana* importin α in complex with the NLS of Tue1. Our structural model is in full agreement with previous experimental importin-α structures supporting that Tue1 is actually imported into the nucleus via a direct interaction with importin-α (**Fig. S1**).

**Figure 1:**
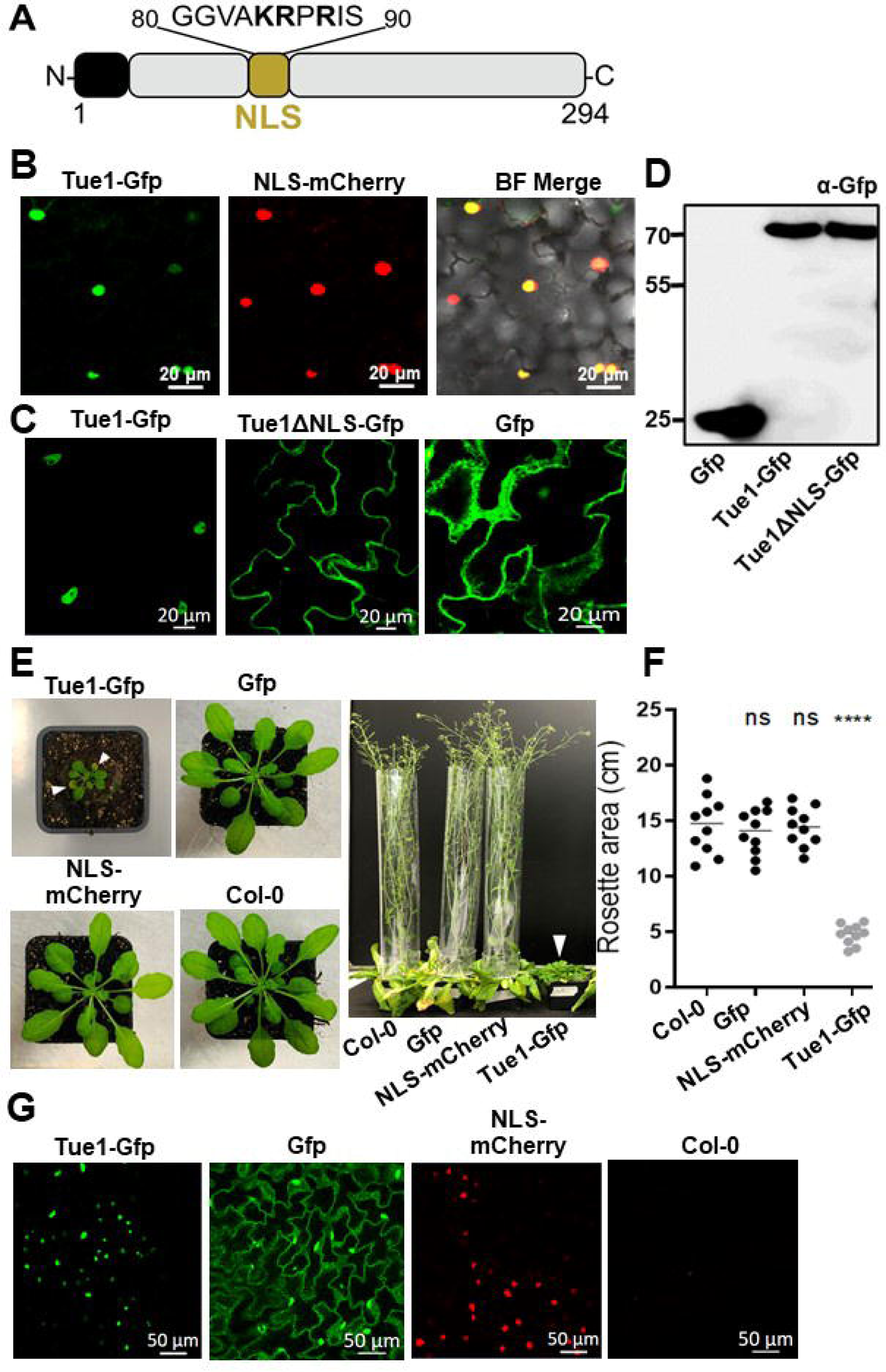
Tue1 localizes to the nucleus of *A. thaliana* and *N. benthamiana*. **A.** Domain architecture of Tue1. The N-terminal signal peptide (SP) and nuclear localization signal (NLS) are indicated as black and yellow boxes, respectively. **B.** Transient expression of Tue1-Gfp in *N. benthamiana* at 3 days post infiltration. The left panel showed the subcellular localization of Tue1 in the nucleus. The middle panel showed the location of the nucleus, visualized by a NLS (At4g19150/N7) marker fused to mCherry. The right panel showed the overlay of GFP and mCherry channels. Yellow spots indicated co-localization of Tue1-Gfp and NLS marker in the nucleus. **C.** Nuclear localization of Tue1-Gfp in *N. benthamiana* in presence and absence of the predicted NLS. Left panel indicated the fluorescence signal of Tue1-Gfp, the middle panel showed signal of the Tue1ΔNLS-Gfp and the right panel indicated free Gfp as a control. NLS truncation in Tue1 leads to cytosolic localization. **D.** Western blot of Tue1-Gfp and Tue1ΔNLS-GFP reveals presence of the full-length fusion proteins. **E.** Macroscopic images of *A. thaliana* lines used in the microscopy experiments. Expression of Tue1-Gfp resulted in chlorosis (white arrowheads) and formation of smaller rosettes as well as delayed in flowering **F.** Quantification of rosette size from the different *A. thaliana* lines showed a significant (p < 0.001) decrease in rosette size in the Tue1-Gfp line. Statistical analysis was done by one-way ANOVA (Bonferroni’s post-test) **G.** Subcellular localization of Tue1-Gfp in transgenic *Arabidopsis thaliana* lines. Microscopic analysis of the stable lines confirmed the nuclear localization of Tue1-Gfp. NLS-mCherry clearly localizes in nuclei, while free GFP localizes both in the cytosol and the nucleus due to its small size.

Importantly, stable over-expression of Tue1-Gfp in *A. thaliana* caused a strong plant growth retardation and delayed flowering, similar to untagged Tue1 (Courville *et al*., 2019), indicating that the fusion protein is functional (**Fig. 1E, F**). Similar to *N. benthamiana*, Tue1-Gfp also showed a clear nuclear localization in *A. thaliana* (**Fig. 1G**). In conclusion, we could show that Tue1 localizes to the plant nucleus via an internal NLS sequence.

### The crystal structure of Tue1 reveals a α-β-α like sandwich architecture

Sequence comparison had not given indications towards the molecular function of Tue1, therefore, we turned to structural analysis. Unfortunately, prediction tools such as AlphaFold2 (Jumper *et al*., 2021) failed to yield reliable results. Therefore, we crystalized the protein to obtain structural insights. Tue1 consists of 294 amino acids with a predicted molecular weight (MW) of 32.6 kDa. We produced Tue1 lacking its signal peptide in *E. coli* and purified the protein using a two-step purification protocol (Zweng *et al*., 2023)(**Fig. S2**). We determined the crystal structure of Tue1 at 1.3 Å resolution, using selenomethionine single-wavelength anomalous dispersion (Se-SAD; **Tab. S1**) as models obtained by AlphaFold2 (Jumper *et al*., 2021) were not of suitable quality for structure-solution by molecular replacement (MR). Amino acids 86 to 257 could be unambiguously modelled into the electron density revealing 9 α-helices and 6 β-strands that fold into a α-β-α like sandwich structure (**Fig. 2A, B**). The first five α-helices form the upper part of the structure that is separated from helices 6-9 by a 6-stranded β-sheet (**Fig. 2B**). The predicted NLS of Tue1 is directly adjacent to helix α1 but could not be resolved in our structure, likely due to flexibility. This supports that the cytosolic localization of the NLS deletion construct (**Fig. 1D**) is not caused by altered protein structure, but due to lack of interaction with the nuclear import machinery (**Fig. S1**).

**Figure 2:**
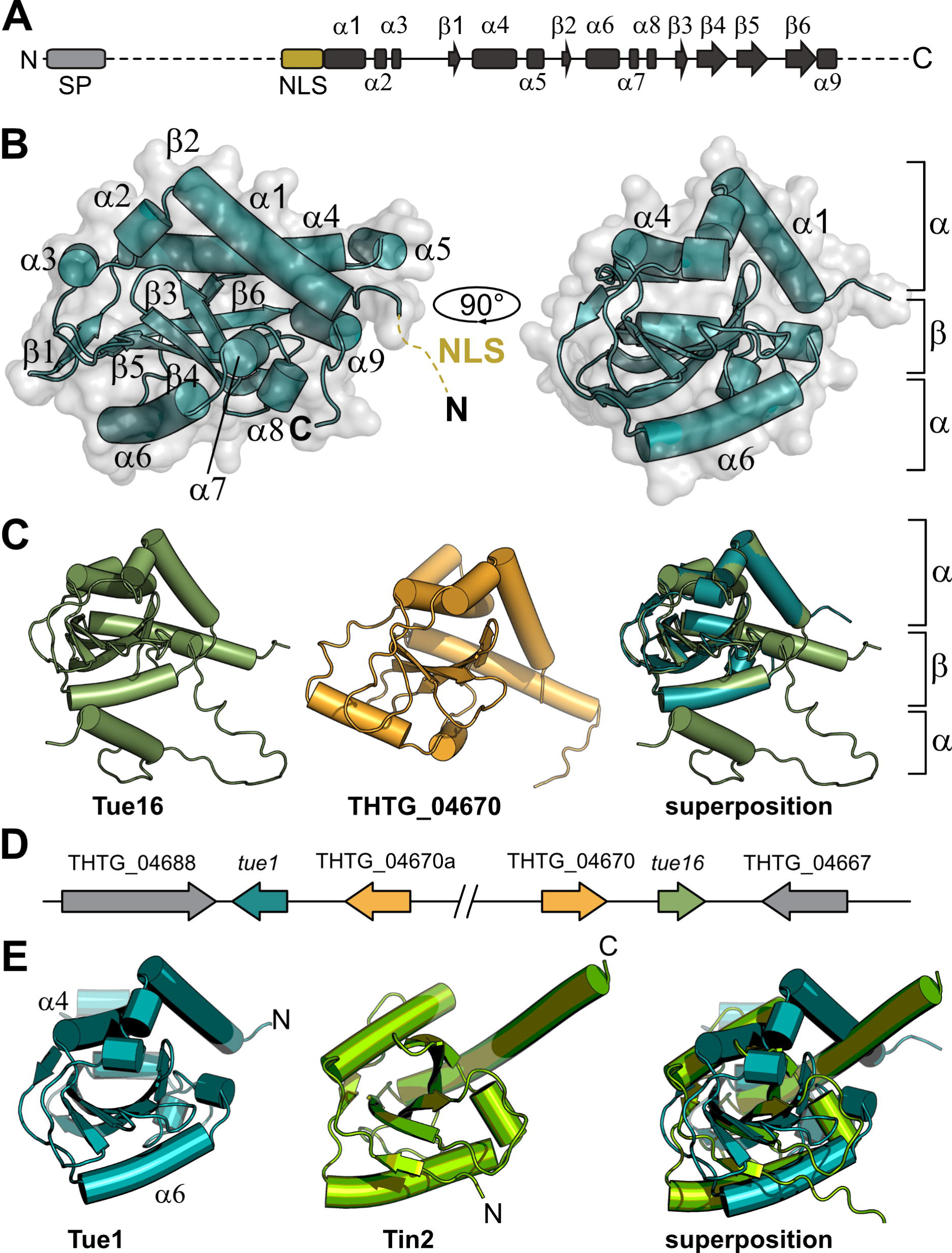
Crystal structure of Tue1. **A.** Domain architecture and secondary structure elements of Tue1. Signal peptide (SP) and nuclear localization sequence (NLS) is displayed in grey and dark green, respectively. Regions not resolved in the electron density are indicated with dashed lines. **B.** Crystal structure of Tue1 displayed as cartoon. The position of the NLS is directly adjacent to α1 and displayed as dashed line. The overall fold resembles an α-β-α sandwich. **C.** Superposition of Tue1, Tue16 (smudge) and THTG_04670 (orange). **D.** Genomic region containing *tue1*, *tue16* and THTG_04670 with the different genes coloured according to the structures.

Interestingly, *tue16*, a second *T. thlaspeos* effector-encoding gene is located on contig 31 in vicinity of *tue1,* and the proteins have a sequence identity of 68.7 % (**Fig. S3**). To compare the two related effector candidates, we predicted the structural model of Tue16 using AlphaFold2 through ColabFold. As expected, Tue16 is structurally highly similar to Tue1 with r.m.s.d.’s of 0.62 (132 Cα) (**Fig. 2C**). As observed before for Tue1, the prediction quality of Tue16 was low due to the lack of available similar sequences in the UniRef database until we used the X-ray structure of Tue1 as query. When inspecting the genomic context, we observed that the whole locus apparently duplicated with *tue1* and *tue16* diverging on the sequence level, while the two copies of another gene, THTG_04670 retained a nucleotide sequence identity of 99.8 % in a 3 kb region of the genomic locus (**Figs. 2D, S3**). We therefore also predicted the structural model of THTG_04670 which has an r.m.s.d. to Tue1 of 2.2 (104 Cα) (**Fig. 2C**). Both, Tue16 and THTG_04670 share the general architecture of several α-helices surrounding a central β-sheet, with the surrounding α-helices being slightly displaced in THTG_04670. In contrast to Tue1 and Tue16, THTG_04670 does not contain a predicted signal peptide (**Fig. S3**) and also shows deviation in the region of the potential NLS. It will be interesting to study, whether THTG_04670 also has virulence activity. In conclusion, determination of the Tue1 crystal structure revealed a α-β-α like sandwich structure that is also shared with another potential effector Tue16 and another protein of yet unknown function, THTG_04670.

### Tue1 is part of an extended structural effector protein family of smut fungi and shares structural homology with intracellular effector proteins of *Ustilago maydis*

With the structural information at hand, we asked if we could identify structural homologs of Tue1 in other plant pathogens. A structural conservation of this fold potentially enables to draw conclusions about the potential function of Tue1 during plant infection. We therefore performed a structural blast using FoldSeek (van Kempen *et al*., 2023) with the crystal structure of Tue1 as query.

We restricted our search to the group of the Ustilaginales as previous searches did not result in significant hits with a convincing structural homology in other organisms. In total, we could identify more than 300 structural homologs of Tue1 from different smut fungal species with Tm-scores ranging from 0.5 to 0.75. High Tm-scores of >0.6 were mostly obtained for proteins from *Pseudozyma flocculosa* and *Sporisorium scitamineum* although the sequence identity was below 10 % in most of the cases. As the structurally related potential effectors of *P. flocculosa* and *Sp. scitamineum* have not yet been functionally characterized, we had a closer look at the candidate with the highest structural similarity from the well-characterized smut fungus *U. maydis*. Tin2 (UMAG_05302) has a Tm-score of 0.54 to Tue1 and shows an overall similar structural architecture (**Fig. 2E**). This effector protein has been thoroughly characterized and acts on the maize protein kinase ZmTTK1, thereby inducing anthocyanin biosynthesis during plant colonization (Tanaka *et al*., 2014). Tin2 is an intracellular effector protein but it does not localize to the nucleus and a closer inspection of the structural elements of Tue1 and Tin2 revealed that the two proteins also show structural differences (**Fig. 2E**). The central β-sheet and spatial positioning of helix α4 and α6 are similar in both proteins, but structural elements decorating the central fold show differences (**Fig. 2E**). Despite Tue1 and Tin2 belonging to the same structural effector family, these structural deviations therefore suggest functional diversification by adapting the core structure. This potentially results in a different interactome during plant colonization of the two hosts of *T. thlaspeos* and *U. maydis*. Interestingly, the sequence related Tin2 homologs from the head smut fungus *Sp. reilianum* have functionally diverged (Tanaka *et al*., 2019). Hence, our structural analysis of Tue1 has revealed that this protein is not unique to *T. thlaspeos* as anticipated from the primary sequence but member of the Tin2-fold family, a structural effector family previously reported in smut fungi (Seong and Krasileva, 2023).

### Tue1 interacts with the jasmonate-ZIM domain protein JAZ10

Functional diversification based on a structural scaffold is a common principle that enables pathogens to evolve target binding while maintaining a robust core scaffold (Derbyshire and Raffaele, 2023). As the structure of Tue1 did not reveal similarities to enzymes or nucleic acid binding proteins, we followed up the hypothesis that Tue1 might target host proteins in the plant nucleus. We therefore aimed to identify potential plant targets in a yeast-two-hybrid (Y2H) analysis. We used a stress-induced cDNA library of *A. thaliana* leaves that were exposed to various biotic and abiotic stresses such as chemical treatments and bacterial inoculation (Matiolli and Melotto, 2018). Tue1 lacking the signal peptide was used as a bait. Co-transformation of a Tue1 expression plasmid with the prey pB42AD plasmid revealed no auto-activation upon analysis of the reporter gene (**Fig. S4A**). This confirms that Tue1 does not possess any elements that might facilitate a direct transcriptional regulation as it has been described e.g. for the activation domain in the b transcription factors *Tt*bW1 and *Tt*bW2 (Frantzeskakis *et al*., 2017). Expression of the full-length Tue1-LexA-DBD was confirmed by Western blotting (**Fig. S4B**).

In total, we identified 140 interaction partner candidates in the screen, for which the respective plasmids were individually transformed to exclude auto-activation and co-transformed with the Tue1 bait vector for validation. Sequencing confirmed in-frame fusions with the DBD, and narrowed the candidate list down to 129 candidates and 110 unique genes (**Supplementary dataset Y2H**). Many of the encoded proteins show functions related to photosynthesis e.g. subunits of the photosystems or RubisCo. Since Tue1 is localized in the nucleus, we filtered the candidates accordingly. Using SUBA5 (Hooper *et al*., 2017), we limited our list to 13 candidates with nuclear localization (**Tab. S6**). Among these, JAZ10 attracted our attention, as we previously noticed that plant hormone-regulated genes are differentially expressed during infection (Courville *et al*., 2019) and JAZ proteins are known targets of several effector proteins interfering with plant defence hormone signalling (Tanaka *et al*., 2015). We therefore focussed on this candidate to validate and characterize the interaction in more detail.

### Tue1 but not Tue16 interacts with JAZ10 proteins from *Ar. hirsuta* and *A. thaliana*

To confirm the interaction of Tue1 with JAZ10 with full-length proteins, we cloned the gene encoding JAZ10 from *A. thaliana* and identified the homologs from the natural host *Ar. hirsuta* in our transcriptome dataset (Courville *et al*., 2019). *Ah*JAZ10 has an amino acid sequence identity of 83 % and contains the conserved CMID, ZIM and Jas motifs (**Fig. S5A**). Targeted Y2H-assay with Tue1 and both JAZ10 proteins form *A. thaliana* and *Ar. hirsuta* (**Fig. S5B**) confirmed that JAZ10 proteins from both plants interact with Tue1 (**Fig. 3A**). In a next step, we aimed to confirm the interaction by bimolecular fluorescence complementation (BiFC) in *N. benthamiana*. Tue1 was fused to the N-terminal half of mVenus and the two JAZ10 proteins were fused to the C-terminal half of mVenus. We used Tue16 as control, as we could previously show that Tue1 and Tue16 have a high structural similarity, share the NLS sequence but show differences on the sequence level (**Fig. 2C**). As positive control, we used a truncated version of the transcription factor MYC3 from *A. thaliana*, a known target of the JAZ10 repressor protein (Fernandez-Calvo *et al*., 2011) that only contained the Jas-binding domain.

**Figure 3:**
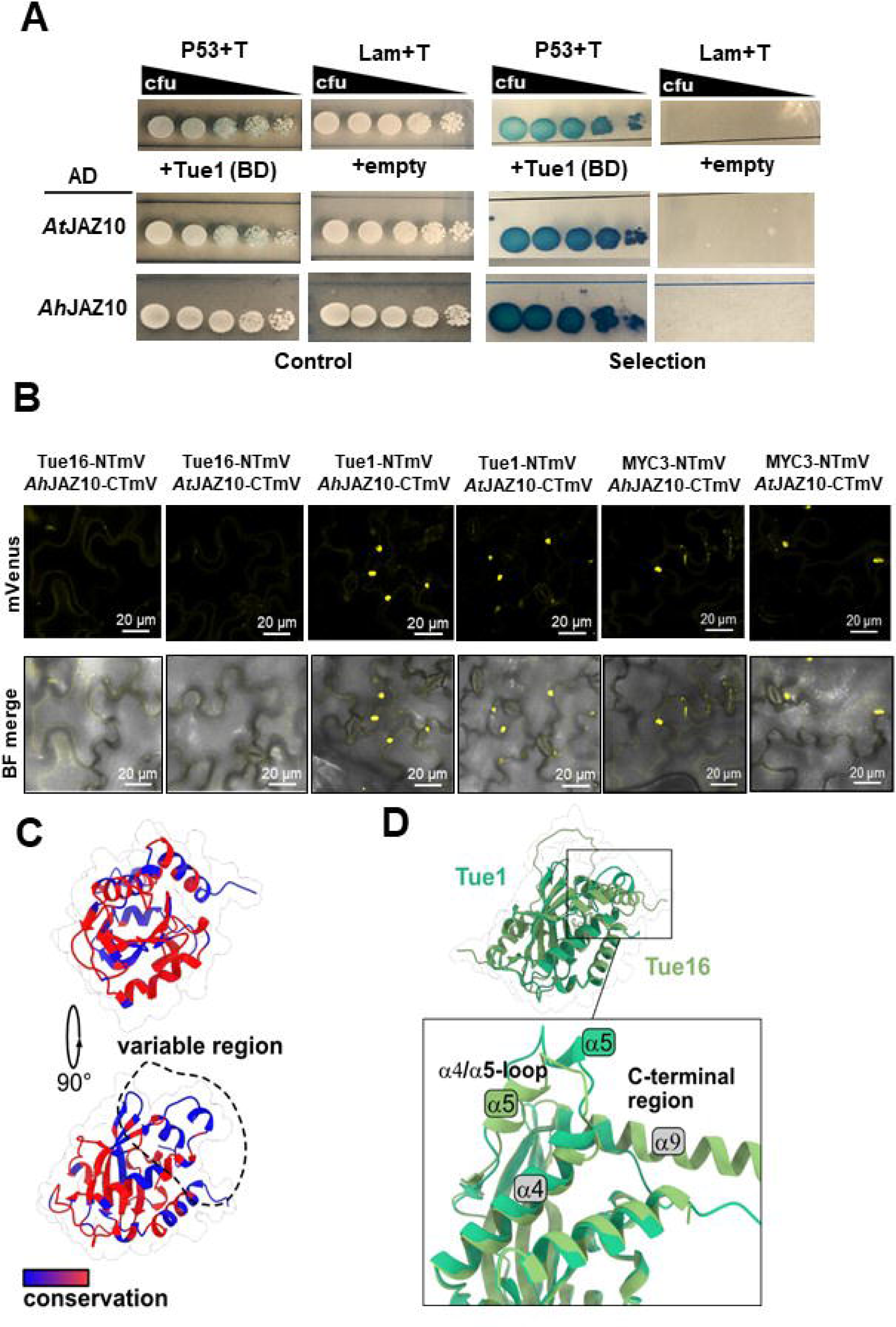
JAZ10 homologs from *A. thaliana* and *Ar. hirsuta* bind Tue1 but not Tue16. Targeted Y2H assay show *in vivo* interaction of both JAZ10 homologs with Tue1. A stringent variant of Y2H assay with X-Gal overlay for expression of a second reporter (LacZ gene) shows comparable results by blue color accumulation with different intensities. Lam (BD) + SV40 T-antigen (AD) as a negative control and P53 (BD) + SV40 T-antigen (AD) as a positive control were shown on the top of each panel. Triple dropout media (SD-His-Trp-Ura) was used for control and quadruple dropout media (SD-His-Trp-Ura-Leu) was used for the selection. **B.** Interaction *in planta* was shown by bimolecular fluorescence complementation (BiFC). mVenus split parts reconstituted and accumulated fluorescence signal in the nucleus upon interaction of fused proteins. Tue16 did not show any fluorescence signal. Fluorophore reconstitution also detected for positive control MYC3. **C.** Based on sequence alignment, Tue1 has a variable region comprising the α4/α5-loop (residues 147-160) and the C-terminal region including α9 (250-257) **D.** The variable region of Tue1 shows a slight structural variance compared to Tue16, which might explain the differences in binding to JAZ10. The inset shows a close-up of the structural regions with the highest differences with secondary structure elements labelled accordingly.

The co-transformation of MYC3 and Tue1 with both JAZ10 homologs showed a clear mVenus fluorescence in the nucleus, while we did not observe any fluorescence when using Tue16 (**Fig. 3B**). Therefore, our BiFC experiments confirmed the interaction between Tue1 and JAZ10, while the closely related paralog Tue16 does not interact with JAZ10 and potentially targets different interaction partners. To investigate the differences between Tue1 and Tue16 that might hint to changes in target specificity or even provide insights about the possible JAZ10 binding site at Tue1, we compared the two structures in more detail. As outlined earlier, the Tue1 and Tue16 are overall highly similar (**Fig. 2C**). Colouring the structure of Tue1 based on the sequence alignment, one region comprising the α4/α5-loop (residues 147-160) and the C-terminal region including α9 (250-257) show a larger degree of sequence variability (**Figs. S3, 3C**). These regions also show a slight structural variance (**Fig. 3D**). Although we cannot rule out that this is a result of the lower confidence of the Tue16 prediction in this region, it might hint to a possible binding interface towards JAZ10.

Next, we aimed to validate the Tue1-JAZ10 interaction *in vitro*. Tue1 and *At*JAZ10 were heterologously produced in *E. coli*. We furthermore produced the Jas-binding domain of MYC3 from *Arabidopsis thaliana,* which was previously used as a positive control in our BiFC experiments (MYC3 5-242). Tue1 and MYC3 were fluorescently labelled and subjected to MST. *At*JAZ10 was titrated in decreasing concentrations starting from 100 µM. We could determine a Kd of 0.43 ± 0.38 µM for the MYC3-JAZ10 interaction, which is in line with a Kd of 5 µM determined previously (Takaoka *et al*., 2022) . The Kd of Tue1 to JAZ10 was lower with 72.53 ± 0.94 µM (**Fig. S6, Tab. S7**). In accordance with our BiFC experiments, no interaction of Tue16 and JAZ10 could be observed (**Fig. S6**). Our MST experiments therefore confirm a direct interaction between Tue1 and JAZ10.

In conclusion, we t Tue16 interact with the Jas domain of JAZ10.

### JAZ10 interacts with the C-terminal region of Tue1

Unfortunately, several attempts to reconstitute the Tue1-JAZ10 complex for structural analysis were unsuccessful and the low confidence of our structural models also prevented complex prediction through AlphaFold2. We therefore decided to follow a structure-guided approach to identify the JAZ10 binding interface at Tue1. Two truncation constructs of *tue1* were generated that code for protein versions either lacking the N-terminal residues 24 to 79 (**Fig. 4A**, Tue1-ΔN) or the C-terminal residues 274 to 294 (**Fig. 4A**, Tue1-ΔC). Both truncations still contained the NLS sequence necessary for nuclear localization and the structural core of the protein containing the Tin2-fold domain (**Fig. 4A**). They were fused to the N-terminal half of mVenus for transient expression in *N. benthamiana*. *At*JAZ10 fused with the C-terminal split mVenus was co-transformed with full length Tue1 and its truncated versions. MYC3 and Tue16 were used as controls. We furthermore included an NLS-mCherry construct in our experiment, not only allowing us to evaluate nuclear co-localization, but also fluorescence signal correlation and quantification. We observed nuclear localization for MYC3, Tue1 and the two truncated Tue1 versions that also correlated with NLS-mCherry fluorescence, indicating that JAZ10 is able to bind all protein versions (**Fig. 4B**). However, we observed a slightly lower fluorescence intensity of co-expressedTue1ΔC and *At*JAZ10 (**Fig. 4B**). We therefore quantified the fluorescence signal intensity for both mCherry and mVenus through mean fluorescence intensity and Pearson coefficient. This analysis unambiguously revealed that, the fluorescence intensity of Tue1, Tue1ΔN and MYC3 co-expressed with *At*JAZ10 correlated with mCherry fluorescence, while signal intensity of Tue1ΔC in combination with JAZ10 decreased significantly (**Fig. 4C, D**) to almost 50 %. The C-terminal residues 274 to 294 also contain helixα9, which is absent in Tue16 (**Fig. 3D**) supporting that JAZ10 is indeed binding to that region of Tue1. To rule out that this is a result of lower protein expression of Tue1ΔC, we performed Western blotting (**Fig. S7**) Indeed, it confirmed that Tue1ΔC fused to the N-terminal half of mVenus is expressed*in planta* in comparable amount as the full-length Tue1-fusion construct (**Fig. S7**).

**Figure 4:**
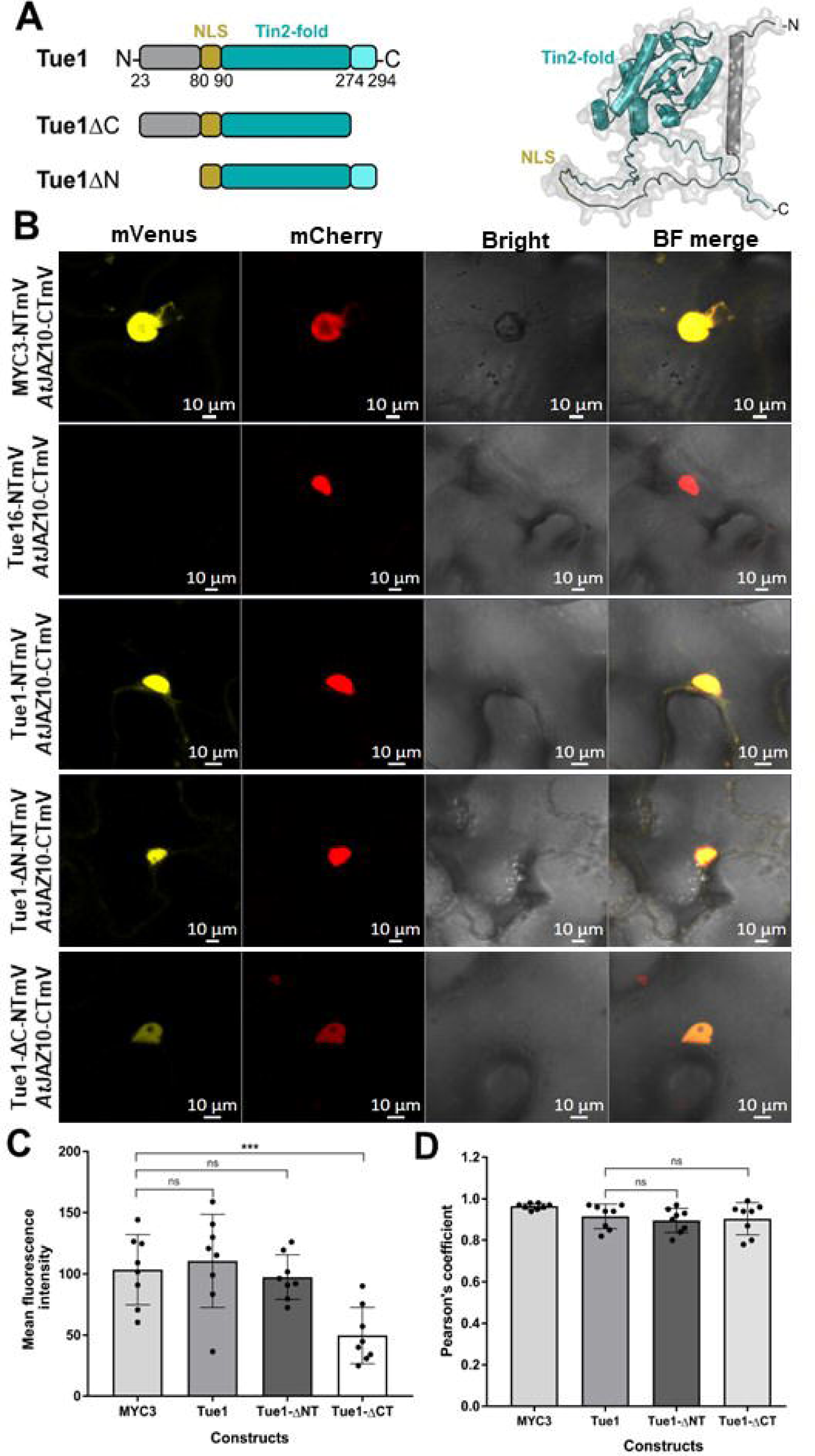
JAZ10 binds to a Tue1 interface that involves the C-terminal residues. **A.** Two truncated constructs of Tue1 were generated that either lacked the N-terminal residues 1-79 (Tue1-ΔN) or the C-terminal residues 274-294 (Tue1-ΔC). Both truncations still contain the NLS sequence necessary for nuclear localization and the structural core of the protein containing the Tin2-fold domain. **B.** The constructs fused with the N-terminal half of mVenus (Tue1-NTmV, Tue1-ΔN-NTmV, Tue1-ΔC-NTmV, MYC3-NTmV) co-expressed with the JAZ10-CTmV in *N. benthamiana*. NLS-mCherry was co-expressed as nuclear marker. **C.** Quantification of fluorescence intensity of mCherry and mVenus by measuring the mean fluorescence intensity and Pearson coefficient. The fluorescence intensity of Tue1, Tue1-ΔN and MYC3 correlated with mCherry while the signal intensity was significantly reduced to 50 % in Tue1-ΔC. **D.** The co-expression of NLS-mCherry construct in our experiment, not only allow us to evaluate nuclear co-localization but also fluorescence signal correlation. Bars indicate mean ±SEM for n=10 technical replicates. ***p<0.05, not significant (ns.), unpaired, two-tailed T-test.

In conclusion, our structure-guided BiFC experiments therefore revealed that JAZ10 likely binds to an interface at Tue1 that involves the C-terminal residues of Tue1 and the lack thereof destabilized the interaction significantly.

## Discussion

Tue1 was previously identified as one of the *Thecaphora*-unique effector candidates that strongly impacted plant development upon overexpression in *A. thaliana* (Courville *et al*., 2019). Here, we performed a detailed analysis of this effector protein providing insights into its molecular function. *In planta*, Tue1 interacts with JAZ10, a repressor of jasmonate signalling. Our structural analysis furthermore revealed that Tue1 is a member of the Tin2-fold family, a previously identified structural family of effector proteins in *U. maydis* (Seong and Krasileva, 2023) with at least 2 members in the *T. thlaspeos* genome. Notably, there is also structural similarity to effectors from other smut fungi, which is not reflected at the sequence level.

### Structural similarity but functional diversity among Tin2-fold smut fungal effectors

Recent advancements in computational methodologies for protein structure prediction, such as AlphaFold (Jumper *et al*., 2021) and RosettaFold (Krishna *et al*., 2024; Mansoor *et al*., 2023), have enabled genome-wide structural predictions of effector proteins. These approaches unveiled that effectors despite being unrelated on the sequence level form structural classes that are often distributed across a variety of host-pathogen systems, examples being RNAse-fold or ToxA-like effectors (Derbyshire and Raffaele, 2023; Seong and Krasileva, 2023). Members of these families have been known earlier but their broad distribution across different pathogens was an astonishing new finding. *U. maydis*, the only smut fungus that was part of this study, harboured only one extended structural family termed Tin2-fold that was not present in any other fungal pathogens (Seong and Krasileva, 2023). Here, we show that the Tin2-fold family is not exclusive to *U. maydis* but can also be found in the distantly related smut *T. thlaspeos* with Tue1 and Tue16 sharing the structural core fold. In *U. maydis*, at least 45 effector proteins have been identified that share the Tin2-fold (Seong and Krasileva, 2023). The majority of these genes are located within gene cluster 19A, a 40 kb genomic segment that is crucial for virulence of *U. maydis* (Brefort *et al*., 2014; Kämper *et al*., 2006). The effector Tin2 also encoded within this segment promotes virulence by sequestering maize protein kinases, thereby altering anthocyanin biosynthesis (Tanaka *et al*., 2014; Tanaka *et al*., 2019). Interestingly, cluster 19A is heavily rearranged in *T. thlaspeos* with all Tin2-fold effector encoding genes being lost from this genomic region (Courville *et al*., 2019). However, our structural analysis now demonstrates that *T. thlaspeos* has not lost all Tin2-fold effectors as Tue1 and Tue 16, both encoded in a small gene cluster, belong to this structural family. Interestingly, Tue16 shares a high sequence identity of 68.8 % with Tue1 and is likely also translocated into the plant nucleus due to a conserved NLS but it does not bind to JAZ10. Our study therefore provides evidence that two closely related Tin2-fold effectors have functionally diversified although the precise function of Tue16 has yet to be clarified. Currently, it is unclear whether such striking functional diversification is a common phenomenon among Tin2-fold effectors that reside within one gene cluster.

Overall, our structural analysis suggests that the Tin2-fold family is shared between all smut fungi where genomic information is available. A detailed analysis has yet to investigate the distribution of Tin2-fold effectors among smut fungi to clarify the evolution and diversification of these effector proteins.

### Tue1 binds to JAZ10 and targets plant jasmonate signalling

We here provide evidence that Tue1 specifically binds to JAZ10, one of the repressors of JA-responsive genes. JA signalling plays a pivotal role in plant defence against fungal pathogens (Antico *et al*., 2012). When plants detect a fungal invasion, they activate the JA pathway, leading to activation of defence gene expression and to the synthesis of defence-related compounds. Perception of JA is mediated by the receptor COI1, an E3 ligase targeting the JAZ repressors for degradation (Li *et al*., 2021; Zhu, 2023). Interestingly, several bacterial effectors are known that specifically target this receptor complex and thereby interfere with JA-dependent defence signalling (Tanaka *et al*., 2015). Coronatine is a bacterial phytotoxin that mimics JA-Ile, activates COI1-mediated proteasomal degradation of the JAZ repressors, and thereby re-opens stomata to enable bacterial entry (Melotto *et al*., 2008; Panchal and Melotto, 2017). HopZ1 and HopX1 from *Pseudomonas syringae* pv. *tomato* DC3000 are enzymes directly target the JAZ proteins as e.g. JAZ2 has been demonstrated to be directly involved in stomata dynamics during a bacterial infection (Gimenez-Ibanez *et al*., 2017). HopZ1 acetylates the JAZ repressors leading to their degradation (Jiang *et al*., 2013); HopX1 is a cysteine protease that directly degrades the JAZ repressor independently of COI1 and the proteasome (Gimenez-Ibanez *et al*., 2014). Interestingly, COI1 and HopZ1 interact with the Jas domain, which is also the binding site for the MYC transcription factors (Takaoka *et al*., 2022), while HopX1 binds to the ZIM domain showing that the different domains can be targeted by effectors. Notably, both bacterial effectors are active enzymes and do not only bind JAZ but rather modulate their targets through acetylation and degradation, respectively (Gimenez-Ibanez *et al*., 2014; Jiang *et al*., 2013). Tue1 also binds to the Jas domain of JAZ10 as demonstrated by our MST experiments. In contrast to the bacterial effectors, we could not detect any hints towards a potential catalytic activity of Tue1. This is in line with previous findings on Tin2-fold effectors as e.g. Tin2 that modulate their targets through blocking the binding sites (Tanaka *et al*., 2014). The affinity of Tue1 towards JAZ10^Jas^ is lower compared to MYC3, which rather supports a model in which Tue1 does not directly competitively inhibit MYC3 binding. Instead, it might stabilize JAZ10 with the help of other factors or it might help transporting JAZ10 into the nucleus similar to MYC2 dependent nuclear import of JAZ1 and JAZ9 (Withers *et al*., 2012) (**Fig. 5**).

**Figure 5:**
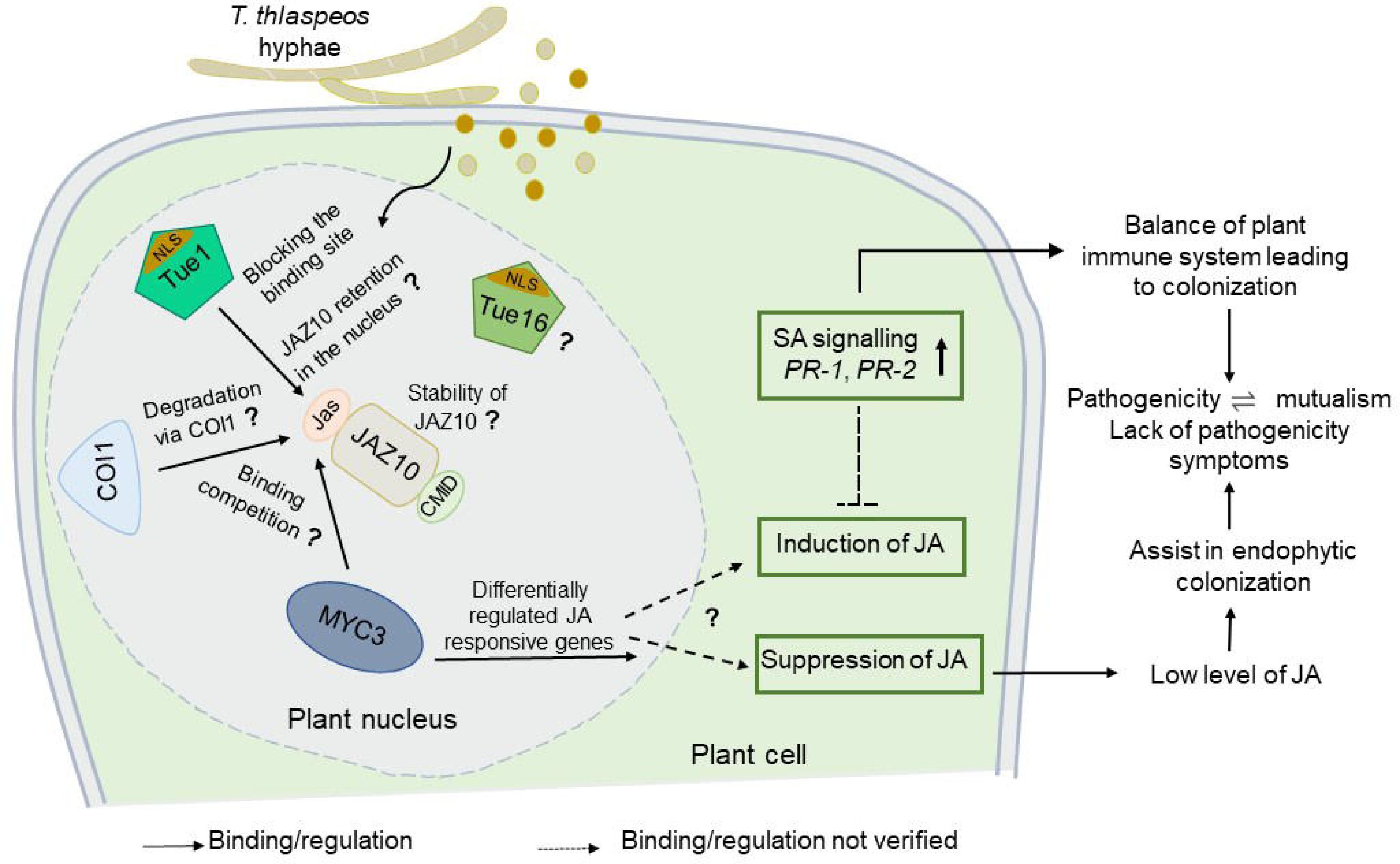
Schematic overview of the possible function of Tue1 during plant infection by *T. thlaspeos*. During *T. thlaspeos* infection, Tue1 and Tue16 are translocated into the plant nucleus. Tue1 binds to the Jas domain of JAZ10, which might serve several purposes: stabilization of JAZ10, retention in the nucleus, or competition with COI1 or MYC3 through blocking the binding site. The role of the related effector Tue16 inside the nucleus is still unclear. Differentially regulated JA-responsive genes and upregulated SA marker genes from transcriptomic data of the *T. thlaspeos* infection demonstrate the suppression of JA pathway. Although plant responses to infection reflect a typical transcriptional change but the pattern of differentially regulated defense genes points towards effective balance. This might limit the excessive fungal proliferation and support colonization without macroscopic symptoms (Courville et al., 2019). Continuous arrow indicates a direct interaction/regulation, dashed arrows indicate an Indirect interaction/regulation not verified.

In contrast to bacteria, fungal effectors targeting JA signalling are less well understood. One example is the ectomycorrhizal interaction between *Laccaria bicolor* and poplar. Here, the small effector protein MiSSP7 interacts with JAZ6 and stabilizes it, resulting in the repression of some JA-responsive genes (Plett *et al*., 2014). Notably, this interaction did not result in differential expression of all JA-responsive genes but only in a subfraction, given that multiple JAZ proteins can redundantly regulate these genes (Pauwels and Goossens, 2011). Comparing the set of differentially regulated genes during *L. bicolor* colonization with our transcriptomic data revealed that there is seemingly not much overlap suggesting a different mechanism for Tue1 than MiSSP7 (**Tab. S8**). This is further supported by a precise analysis of the function of MiSSP7 that strengthens the binding of JAZ6 to MYC2.1 and antagonizes JAZ6 oligomerization (Daguerre *et al*., 2020), which is likely different from the mechanism of Tue1. Many nuclear localized effectors from different pathogens and specifically those that target JAZ family showed their role in exploiting nuclear processes such as hormonal pathways, RNA processing and transcriptional regulation (Harris *et al*., 2023). Instead of an induction, differential regulation of JA responsive genes as well as upregulated SA marker genes from transcriptomic data of the *T. thlaspeos* infection (Courville *et al*., 2019) rather indicate the suppression of JA pathway **(Fig. 5).** *T. thlaspeos* growth in the host plant could benefit from low JA levels similar to an elevated colonization trend described in a rice species colonized by an endophyte *Azoarcus olearius* (Chen *et al*., 2019). Therefore, one hypothesis could be that this is an adaptation to the extended biotrophic phase of the perennial pathogen or limitation to the vasculature, which avoids intensive contact of fungal hyphae with plant cells as found for intracellular hyphae of *U. maydis.* This effective balance in plant defence responses could serve to maintain the symptomless fungal growth and eventually completion of fungal life cycle. Additionally, during the biotrophic phase, Tue1 is not the only effector, so that in the transcriptome, we cannot pinpoint the effect of the Tue1-JAZ10 interaction directly. Further research is therefore required to fully understand the functional relevance of the Tue1-JAZ10 interaction and delineate the precise molecular mechanism of this effector protein. In conclusion, we here provide the first mechanistic insights into an effector protein from the smut fungus *T. thlaspeos*, which supports plant infection by directly binding to JAZ10 of *A. thaliana* and *Ar. hirsuta*. Our structural analysis furthermore suggests the expansion of Tin2-fold effectors to even distantly related smut fungi such as *T. thlaspeos*, providing evidence for an evolutionary conservation of this family.

## Acknowledgements

We thank Michael Feldbrügge and Kerstin Schipper for critical reading and fruitful discussions on the manuscript. We are grateful to Maeli Melotto for providing the cDNA library and support. The Center for Structural Studies is funded by the Deutsche Forschungsgemeinschaft (DFG Grant number 417919780, INST 208/740-1 FUGG and INST 208/868-1 FUGG to S.H.J.S.). This work was supported by the Deutsche Forschungsgemeinschaft (DFG, German Research Foundation) under Germanýs Excellence Strategy – EXC 2048/1 (to F.A. and V.G.), SFB1535 -Project ID 458090666 (to F. A. and S. H.J. S.) and GRK2466 – Project ID 391465903 (to V.G. and G.M.R.). S Gul was supported by a doctoral fellowship of the Schlumberger foundation faculty for the future program. The funders did not have role in experimental planning, data analysis and manuscript preparation.

## Author contributions

VG, FA: conceptualization; FA: formal analysis; SuG, GMR, StG, NH: investigation; SHJS: resources; NH: data curation; SuG, GMR, FA, VG: writing - original draft; FA, VG: writing - review & editing; SuG, GMR, FA: visualization; FA, VG: supervision; SHJS, FA, VG: funding acquisition

## Data availability

All data are contained within this manuscript. The crystal structure of Tue1 was deposited at the PDB (PDB-ID: 9FPM).

